# Affinity of rhodopsin to raft enables the aligned oligomer formation from dimers: Coarse-grained molecular dynamics simulation of disk membranes

**DOI:** 10.1101/850966

**Authors:** Yukito Kaneshige, Fumio Hayashi, Kenichi Morigaki, Yasushi Tanimoto, Hayato Yamashita, Masashi Fuji, Akinori Awazu

## Abstract

The visual photopigment protein rhodopsin (Rh) is a typical G protein-coupled receptor (GPCR) that initiates the phototransduction cascade in retinal disk membrane of rod-photoreceptor cells. Rh molecule has a tendency to form dimer, and the dimer tends to form rows, which is suggested to heighten phototransduction efficiency in single-photon regime. In addition, the dimerization confers Rh an affinity for lipid raft, i.e. raftophilicity. However, the mechanism by which Rh-dimer raftophilicity contributes to the organization of the higher order structure remains unknown. In this study, we performed coarse-grained molecular dynamics simulations of a disk membrane model containing unsaturated lipids, saturated lipids with cholesterol, and Rh-dimers. We described the Rh-dimers by two-dimensional particle populations where the palmitoyl moieties of each Rh exhibits raftophilicity. We simulated the structuring of Rh in a disk for two types of Rh-dimer, i.e., the most and second most stable Rh dimers, which exposes the raftophilic regions at the dimerization-interface (H1/H8 dimer) and two edges away from the interface (H4/H5 dimer), respectively. Our simulations revealed that only the H1/H8 dimer could form a row structure. A small number of raftophilic lipids recruited to and intercalated in a narrow space between H1/H8 dimers stabilize the side-by-side interaction between dimers in a row. Our results implicate that the nano-sized lipid raft domains act as a “glue” to organize the long row structures of Rh-dimers.

## INTRODUCTION

The visual pigment rhodopsin (Rh) initiates the phototransduction cascade in vertebrate disk membranes of rod photoreceptor cells. Rh is a prototypical seven-transmembrane G protein-coupled receptor (GPCR) and is highly concentrated in the disk membrane, occupying approximately 30% of the total disk membrane area [1,2].

Similar to many other GPCRs, Rh is doubly palmitoylated (C16:0) at the C-terminus of the juxta-membrane eighth-helix (H8) [3]. The tandem palmitoyls are known to be a robust raft-targeting signal for membrane proteins [4–9]. However, despite having two palmitoyls, Rh prefers polyunsaturated phospholipids because of its rough intramembrane surface, which is similar to other membrane proteins [10,11]. Thus, rhodopsin is inherently a non-raftophilic (raftophobic) membrane protein. Nevertheless, dimerization, which is stabilized by the binding of the cognate G protein transducin, confers a high lipid raft affinity (raftophilicity) for Rh, i.e., the di-palmitoyl modification at the C-terminus of H8 is prerequisite for the dimerization-dependent raftophilicity of Rh [12]. Single- and semi-multimolecular observations on rhodopsin dynamics in retinal disk have revealed that the oligomerization-induced raftophilicity of Rh promotes spontaneous formation of raftophilic Rh-clusters [13].

The organization and dynamics of Rh in the disk membrane have long been debated. There have been contradictory views regarding Rh, i.e., freely diffusing monomeric Rh [11,13,14], the static and highly ordered rows of Rh dimer [15–18], and multi-stage oligomeric states in dynamic equilibrium [19]. A recent single-molecule and semi-multimolecular study of Rh dynamics revealed that Rh exists in a dynamic equilibrium between three diffusive states, presumably ascribable to the monomer, dimer, and 100 nm-order short-lived clusters of Rh [13]. Accumulating evidence based on biochemical analyses, atomic force microscopy (AFM) and cryo-electron tomography implies that Rh tends to form rows of dimers in the disk membranes [15–18]. Particle-based simulation studies suggest an essential role of the row structure of Rh for efficient and stable signal amplification. It is hypothesized that the single-photon bleached monomeric Rh can activate a specified number of G protein transducin pre-associating with the row structure of Rh dimers [17,20]. Therefore, we propose that the short-lived meso-sized cluster of Rh is a paracrystalline array of Rh dimers.

The mechanism by which Rh-dimers form a regular row structure remains unknown. MARTINI force field is a coarse-grain force field used for molecular dynamics simulations of biomolecular systems. Recent free energy estimations using MARTINI force field suggest two types of Rh-dimer structures are stable, the H1/H8 dimer and the H4/H5 dimer [21]. The H1/H8 dimer is formed when the first alpha-helix of one Rh molecule and the eighth helix of another come in contact with each other (Fig 1a). The H4/H5 dimer is formed between transmembrane helices 4 and 5. While both Rh-dimers appear to be stable, it is unclear which dimer is typical in disk membranes. Thus, in this study, we proposed a coarse-grained molecular dynamics model of disk membranes described by two-dimensional (2-D) particle populations consisted of unsaturated lipids, raftophilic saturated lipids, and one of the two types of Rh-dimers.

**Fig 1.**
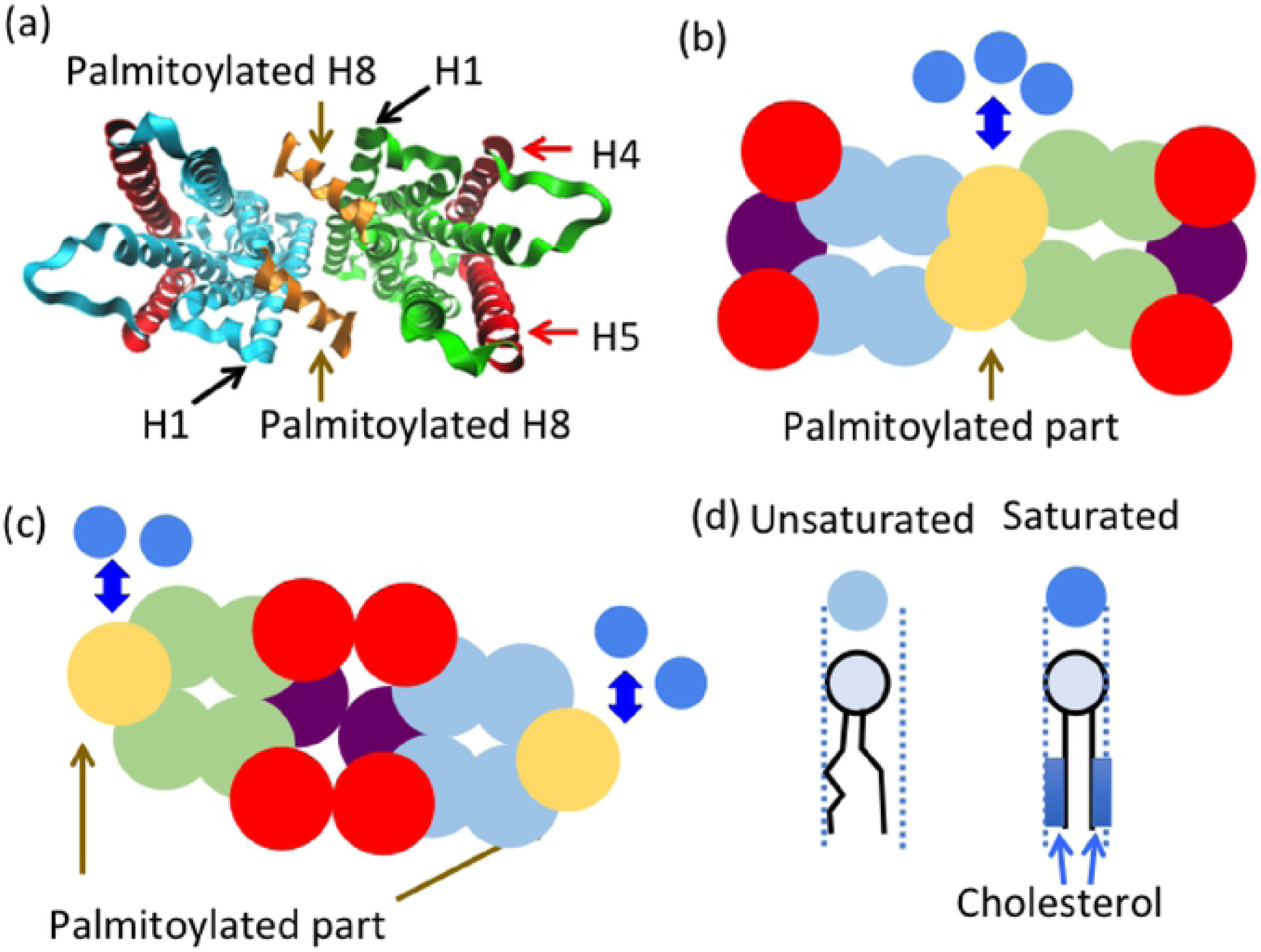
Models of rhodopsin (Rh)-dimers and unsaturated and saturated lipids. (a) Illustration of the three-dimensional structure of Rh based on X-ray crystal structure analysis of the deactivated form of the H1/H8 dimer (PDB ID:3CAP) [22]. The ribbon model of H1/H8 dimer viewed from the eighth helix (H8) side. The yellow alpha-helix indicates H8 in each Rh-monomer, which was assumed to be palmitoylated. The red alpha-helices indicate H4 and H5. (b–c) Basic two-dimensional (2-D) structure models. (b) H1/H8 dimer 2-D structure constructed based on the Rh-dimer contour profile in (a). (c) H4/H5 dimer 2-D structure. The yellow particles indicate regions of H8 assumed to be raftophilic. The red particles are assumed to be parts of H4 and H5. The other particles (blue or green) represent other regions of the Rh dimer. (d) Schematic illustrations of unsaturated and saturated lipids (lower figures) and their representative 2-D particle models (upper figures). The affinity between particles of saturated lipids and the yellow particles in the Rh-dimer 2-D model was assumed (see panels b and c). The tails of each saturated lipid and cholesterol were assumed to frequently associate with each other, making the saturated lipids rigid compared to the unsaturated lipids.

We compared the simulation results among the disk membrane models of only H1/H8 dimers and H4/H5 dimers. The raftophilic acyl chains of the eighth alpha helix were located at the center of each H1/H8 dimer [12] while the acyl chains of the H4/H5 dimer were located at the two outer edges. Our simulation suggested the shapes of Rh-oligomers differed among the models with only the model of H1/H8 dimers predicting that appropriate raftophilicity was able to form ordered row structures of Rh-dimers. Each pair of Rh dimers was connected due to raftophilic saturated lipid domains that existed within their raftophilic regions.

## Model and Methods

### Model concept: Coarse-grained model of Rh and lipids in retinal disk membranes

We constructed a coarse-grained molecular dynamics model of retinal disk membrane to investigate the mechanism of row structure formation by Rh-dimers. Recent studies have suggested that approximately 70–80% of Rh molecules form dimers in the disk membrane [13,18,19]. Furthermore, Rh-dimers are known to locate in the disk membranes where saturated and unsaturated lipids are essential components [13,18]. Therefore, for simplicity we focused on the dynamics of these two types of lipids and Rh-dimers in the current model in order to advance our insight into the mechanisms of row structure formation by Rh-dimers.

The disk membrane is a closed lipid bilayer membrane incorporating a tremendous amount of Rh. The dominant movement of Rh and lipids is 2-D diffusion along the membrane plane. The lipid-lipid and lipid-Rh positional exchanges along the membrane plane are the predominant limiting processes. Again for simplicity, we constructed the disk membrane model using a complete 2-D particles system that described the structures and dynamics of lipids in one monolayer and Rh-dimers on the plane using circular particles (Fig 1). the influences of water and lipids in the opposing disk membrane monolayer were modeled as drag forces and noise.

The 2-D structure model of each Rh-dimer was constructed as a 2-D elastic network using 16 circular particles. For the H1/H8 dimer model, the particles occupied positions to imitate the contour profile of the basic Rh-dimer structure viewed from the H8 side based on X-ray crystal structure analysis of the deactivated form of Rh-dimers (PDB ID: 3CAP) [22] (Figs 1a–b and S1a). The radii of the particles were assumed to be similar to those of alpha helices and the nearby particles were connected by springs where the natural length of each spring connecting two particles was equal to the distance between them according to the basic 2-D Rh-dimer structure model (S1b Fig and S1 Table). As there are no reference structures based on experiments such as X-ray crystal structure analysis, the H4/H5-dimer model was constructed by exchanging the positions of the particles on the left half side of the H1/H8 dimer model corresponding to one Rh-monomer with those on the opposite side (Figs 1c and S1c–d and S2 Table). We note that the results presented in this paper were independent of the detailed shape of the 2-D model of Rh-dimers and were qualitatively unchanged.

The structures and physical properties of the unsaturated and saturated lipids in the 2-D model were as described. Both types of lipids were assumed as circular particles. The radii of the circular particles were assumed as those when lipids are approximated as cylinders (Fig 1d). No specific attractive interactions were assumed among the lipids. Notably, disk membranes are known to contain sufficient concentrations of the cholesterol (11 molar %) compared to that of saturated lipids and the frequent interactions of cholesterols with saturated lipids are expected to make the saturated lipids rigid and raftophilic compared to that of unsaturated lipids [1]. Therefore, we assumed the cholesterols were always associated with saturated lipids and the lipids were always more rigid than unsaturated lipids. These assumptions resulted in the saturated lipid membranes being more rigid and less elastic than that of unsaturated lipid membranes, which is consistent with experimental findings [23]. Furthermore, as mentioned below, the simulations under these assumptions demonstrated the following results, i) the diffusion of saturated lipids was slower than that of unsaturated lipids and ii) the saturated lipid domains that may correspond to the ordered lipid raft domains appeared through the phase separation between saturated and unsaturated lipids (see the “Parameters for simulations” section in the Results). These results were consistent with previously reported experimental findings [21,24,25].

Additionally, two H8 helices located near the center of the H1/H8 dimer and near the edges of the H4/H5 dimer were expected to be partially raftophilic since acyl chains in H8 are known to be palmitoylated [4,12,26]. Therefore, in the models of these dimers, we assumed an affinity between saturated lipids and the tip of H8 (yellow particles in Fig 1c), except for some modified models as detailed below.

### Model implementation

In the model used in this study, we described the central region of the disk membrane to consist of Rh and two types of lipids using 2-D particles moving on a 2-D plane. Since the lipids and Rh would be subjected to water and lipids in the opposing monolayer of the membrane, we assumed that the motion of each particle, the lipids, or parts of Rh obeyed overdamped Langevin equation, as follows:

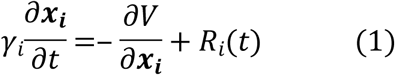

where ***x***_***i***_ = (*x*_*i*_, *y*_*i*_) was the position of the *i*-th particle and *V* indicated potential of forces working on particles. The *γ*_*i*_ and *R*_*i*_(*t*) indicated the coefficient of drag force and the random force working on *i*-th particle by water and lipids in the opposite monolayer, respectively. The term *R*_*i*_(*t*) was given as Gaussian white noise and satisfied ⟨*R*_*i*_(*t*)⟩ = 0, and ⟨*R*_*i*_(*t*)*R*_*j*_(*s*)⟩ = 2*γ*_*i*_*k*_*B*_*Tδ*_*ij*_*δ*(*t* − *s*) where *k*_*B*_ was the Boltzmann constant, *T* was the temperature, *δ*_*ij*_ indicated the Kronecker delta, and *δ* () indicated the Dirac delta function.

The first term of the right-hand side of equation (1) indicated interactions among particles provided by the potential of the system *V* according to the following equation:

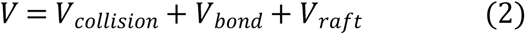

where *V*_*collision*_ was the potential of the excluded volume effects among the particles, *V*_*bond*_ was the interaction potential among the particles forming Rh-dimers to sustain the shape of each dimer, and *V*_*raft*_ was the affinity for the interactions between Rh dimers and saturated lipids.

The potential of the excluded volume effects among particles (*V*_*collision*_) was denoted by the following:

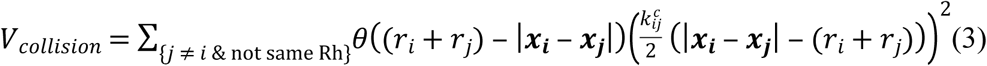

where 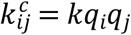 was the elastic constant when the *i*-th and *j*-th particles contacted each other, *q*_*i*_ indicated the nondimensional parameter of rigidity of the *i*-th particle, *r*_*i*_ indicated the radius of the *i*-th particle, and *θ* was the Heaviside step function defined by the following:

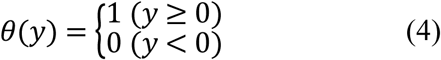

We assumed *q*_*i*_ depended on the type of particles, where *q*_*i*_ of saturated lipids was assumed to be appropriately larger than those of unsaturated lipids. In this case, the lipid type-dependent diffusion constant and the saturated-unsaturated lipid phase separations were consistently obtained based on recent experiments as described below in the Results section and as previously described [21,24,25]. The variables *r*_*h*_ and *r*_*l*_ were the radii of cylinders, which approximated an alpha-helix of Rh and the lipids in the disk membrane, respectively. We assumed *r*_*i*_ = *r*_*h*_ when the *i*-th particle was a component of Rh and *r*_*i*_ = *r*_*l*_ when the *i*-th particle was a lipid. The term Σ indicated the sum of the *i*-th and *j*-th particles that did not belong to the same Rh-dimer.

The interaction potential among the particles forming Rh-dimers was defined as *V*_*bond*_, and was calculated as follows:

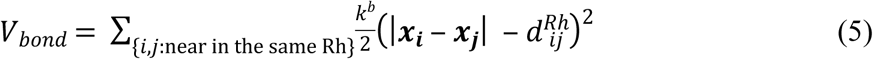

Where 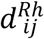 was the distance between the *i*-th and *j*-th particles of the basic 2-D Rh-dimer structure (S1 Fig and S1 and S2 Tables) and *k*^*b*^ was the elastic constant for sustaining the shape of the basic structure of each Rh-dimer. The Σ in this equation indicated the sum of the *i*-th and *j*-th particles that belonged to the same Rh-dimer and that were in close proximity to each other (S1 Fig).

The interaction potential among the palmitoylated H8s of the Rh dimers and the saturated lipids was indicated as *V*_*Raft*_ and was calculated as follows:

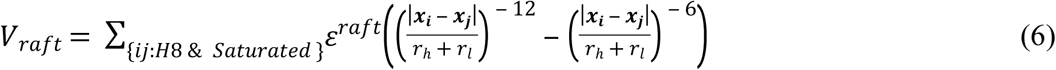

where *ε*^*raft*^ indicated the affinity between a palmitoylated H8 and a saturated lipid. The S indicated the sum of particles pairs of a particle occupied in H8 of Rh and a saturated lipid (Fig 1).

### Simulation parameters

As far as possible, the parameters of the model employed in the simulations of the current study were based on experimental findings. The parameters included radii of lipids (*r*_*l*_) and alpha helix (*r*_*h*_) of Rh and were assumed to be 0.43 nm and 0.88 nm, respectively. To simulate molecular dynamics similar to the situation at the central region of the disk membrane, we used a 40 nm × 40 nm square box with periodic boundary conditions, which included 14 Rh-dimers (14 × 2 × 8 = 224 helices), 156 saturated lipids, and 1,704 unsaturated lipids as entire the simulation space. The area occupancy rates of the Rh-dimer, saturated lipids, and unsaturated lipids in the present model were given as approximately 27%, 5%, and 58%, respectively, which matched the rates for the entire disk membrane obtained from previous experiments [1,2].

The parameters for the excluded volume effect of each particle (*k* and *q*_*i*_) were assumed according to *k* / *k*_*B*_*T* = 7.5 (*nm* ^− 2^), *q*_*i*_ = 1.5 for unsaturated lipids, *q*_*i*_ = 4 for saturated lipids, and *q*_*i*_ = 9 for particles in Rh. To sufficiently sustain the Rh-dimer structure, we assigned *k*^*b*^ / *k*_*B*_*T*∼18870 (*nm* ^− 2^). The drag coefficient *γ*_*i*_ was assumed at *γ*_*i*_ = 6*πηr*_*i*_. In the current model, *η* was expected to be relatively large compared to cytoplasm as we did not know the precise value from previous experiments because of the influences of the lipids at the opposing lipid bilayer. Thus, we assumed *η*/ *k*_*B*_*T* = 6 × 10 ^− 6^(*nm* ^− 3^*s*) in the current arguments, while *η*/ *k*_*B*_*T*∼1.55 × 10 ^− 7^ (*nm* ^− 3^*s*) is estimated in cytoplasm [27]. In this case, the additional simulation of the model using these values for the parameters without any affinity among the lipids and Rh-dimer (such a dimer was termed a raftophobic H1/H8 dimer and is defined in the next section), we obtained diffusion coefficients for unsaturated lipids (in an unsaturated lipid domain), saturated lipids (in a saturated lipid domain), and Rh-dimer (in an unsaturated lipid domain) of approximately 27 (*μm*^2^/*s*), 3.0 (*μm*^2^/*s*), and 0.5 (*μm*^2^/*s*), respectively. These values were estimated by mean square displacement of the particles (S2 Fig and S3 Table). The diffusion coefficients were consistent with those observed in recent *in vitro* experiments using artificial model membranes [11,13,28,29].

The parameter used to estimate the affinity between the particle at H8 of the Rh-dimer and the saturated lipid was assumed according to *ε*^*raft*^/*k*_*B*_*T* = 62.5, which was based on the results of the following additional simulation of a model membrane containing only one Rh-dimer and an equal number of saturated and unsaturated lipids with the abovementioned parameters (S3 Fig). Using this simulation, we obtained the probability of contact between the Rh-dimer and saturated lipids as approximately 0.627. This result seemed consistent with our *in vitro* study using artificial lipid bilayers containing saturated and unsaturated lipid domain where approximately 62% of the Rh-dimers with H8 palmitoylation were found to have saturated lipid domains at equilibrium (Tanimoto, et al., submitted). However, we note that the detailed values *ε*^*raft*^/*k*_*B*_*T* were not essential and similar contact probability values could be obtained when *ε*^*raft*^/*k*_*B*_*T* ≫ 1.

### Analysis methodology

To estimate the degree of order of the spatial distribution of Rh-dimers, we defined the degree of row structure *S*(*t*) as follows:

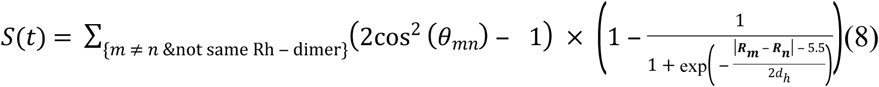

where the position of the center of mass of the *m*-th Rh-dimer (***R***_***m***_ and *θ*_*mn*_) and the angle between the orientation of the *m*-th and *n*-th Rh dimers as shown in S4 Fig. The contribution of *m*-th and *n*-th Rh-dimers to *S*(*t*) was large when the centers of masses were close to each other and when they were facing the same direction. The value of cos (*θ*_*mn*_) was determined from the inner product between the vector from one Rh-Rh interface particle to another of the *m*-th Rh-dimer and that of the *n*-th Rh-dimer (S4 Fig), where 2(cos^2^ (*θ*_*mn*_) − 1) was used as the degree of plane orientation in liquid crystal [30]. Since the distance between Rh-dimers observed in cryo-electron tomography is approximately 5.5 *nm* [4], we assumed *S*(*t*) drastically decreased when the distance between two Rh-dimers was greater than 5.5 *nm*. Ideally, *S*(*t*) = 0 when Rh-dimers were distributed randomly.

## Results

### H1/H8 dimers could form row structures of Rh-dimers by their raftophilicities

The model simulations of disk membrane containing H1/H8 dimers were performed in the current study. We assumed all molecules were randomly distributed under the initial conditions (Fig 2a). Snapshots of the simulations after extended periods of time from the initial condition revealed that the model with H1/H8 dimers exhibited some row structures of Rh-dimers (Fig 2b), consistent with findings observed in AFM and cryo-electron tomography [15–18]. On the other hand, no Rh-dimer row structures were observed in models using raftophobic dimers (Fig 2c), where Raftophobic H1/H8 dimers indicates H1/H8 dimers but assumed *ε*^*raft*^ = 0. According to the measurement of *S*(*t*) and their averages over 10 simulations (⟨*S*(*t*)⟩), the model using H1/H8 dimers exhibited larger *S*(*t*) and ⟨*S*(*t*)⟩ values and greater averages after extended time from the initial conditions compared to those using raftophobic dimers (Fig 3).

**Fig 2.**
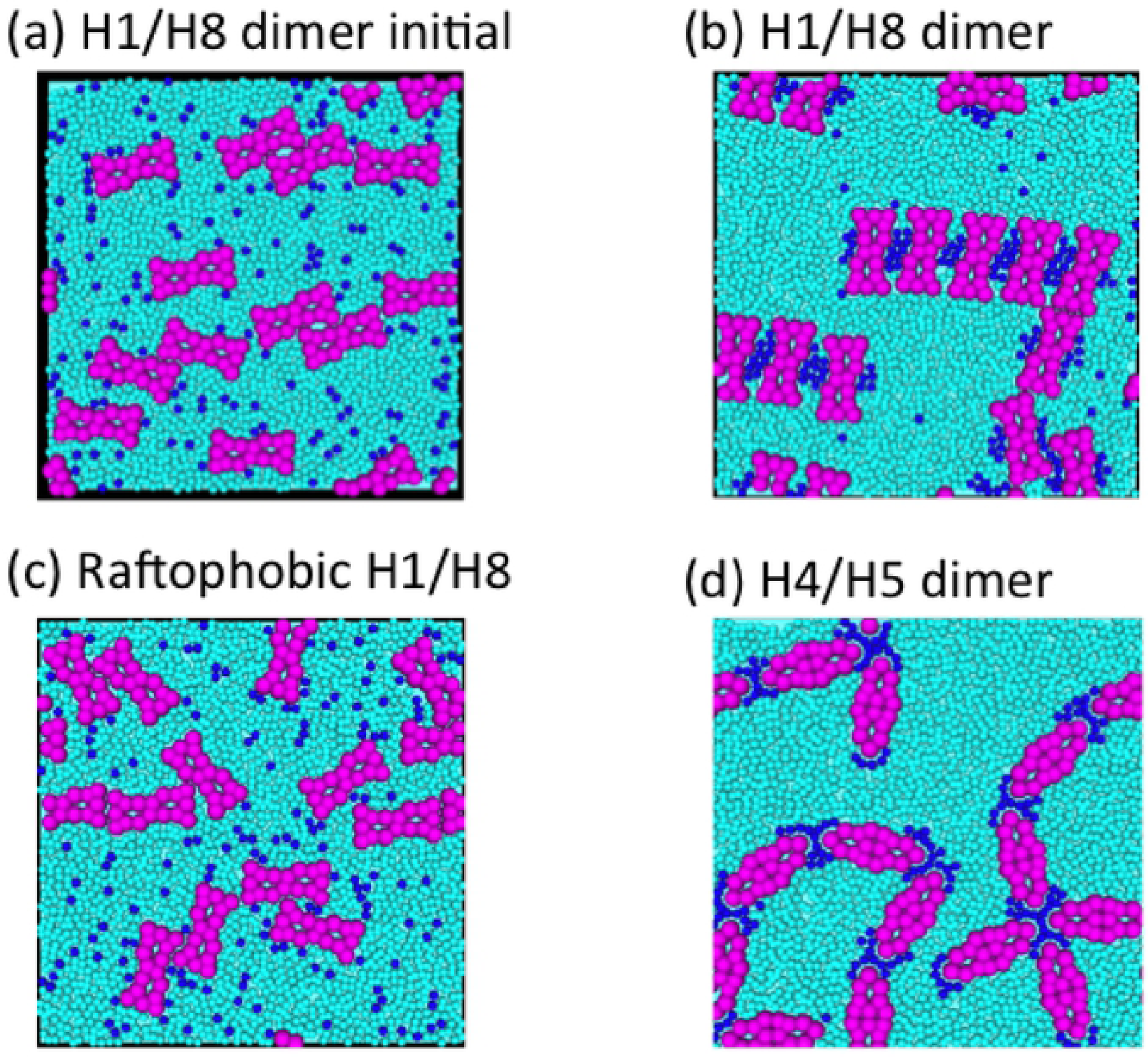
Typical snapshots of rhodopsin (Rh) and lipid distribution obtained from simulations using four models of retinal disk membrane containing Rh H1/H8 dimers and H4/H5 dimers. Snapshots of Rh and lipid configurations at (a) time (t) = 0 s (initial state) and t = 200 µs for models (b) with H1/H8 dimers, and (c) with raftophobic H1/H8 dimers. Raftophobic H1/H8 dimers indicates H1/H8 dimers assumed *ε*^*raft*^ = 0. (d) Snapshots of Rh and lipid configurations at t = 200 µs for models with H4/H5 dimers. Large magenta particle populations indicate Rh proteins. The blue and cyan particles indicate saturated and unsaturated lipids, respectively.

**Fig 3.**
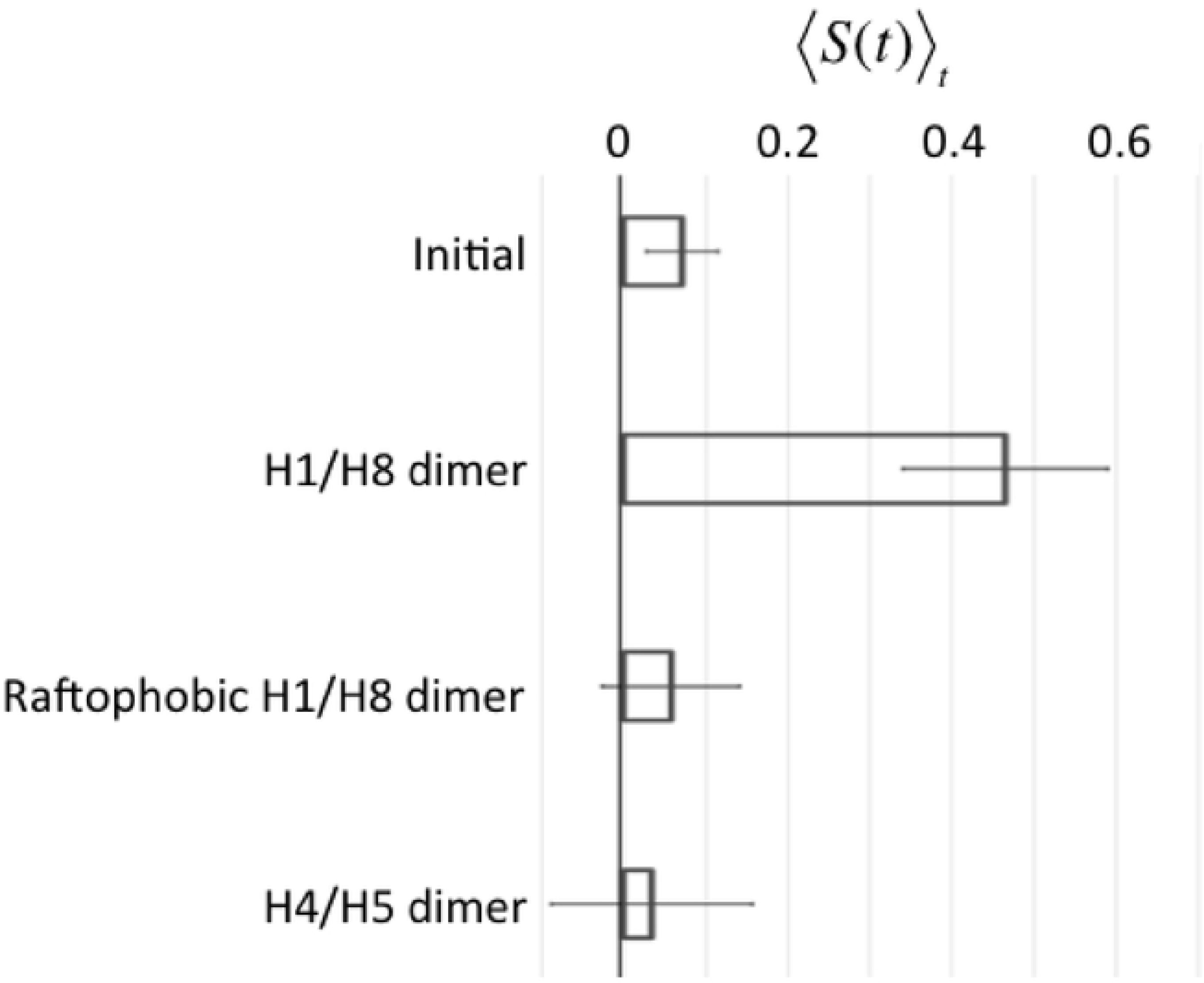
Evaluation of degrees of order of spatial distribution of rhodopsin (Rh) dimers. Averages of ⟨*S*(*t*)⟩_*t*_ over ten simulations, ⟨*S*(*t*)⟩, for each model with a different Rh dimer as indicated. Error bars indicate standard deviations. The “Initial” values were obtained for ten initial conditions for which the configurations of molecules were random. The ⟨*S*(*t*)⟩_*t*_ for each model of dimer indicates the time for the average degree of row structure by the Rh-dimers, *S*(*t*), ranging between 180 and 200 µs.

### H4/H5 dimers accumulated but failed to form row structures of rhodopsin (Rh)-dimers

The simulation of the model containing H4/H5 dimers showed Rh-dimer accumulation with some branching structures, but no row structures were observed (Figs 2d and 3). In models using either dimers, H1/H8 or H4/H5, associations among the Rh-dimers occurred through the small regions of saturated lipid accumulation near the areas corresponding to raftophilic H8s of each Rh-dimer. Notably, for H1/H8 dimers, two dimers associated with each other when the saturated lipid domains formed at the regions between two the central portions of these dimers. Each domain connected only to the central regions of two Rh dimers that corresponded to their raftophilic H8s. This fact enabled the H1/H8 dimers to form long rows of Rh-dimer structures (Figs 2b and 4a). In contrast, H4/H5 dimers could associate with more than two dimers through the domain of saturated lipids that formed at the edges of each dimer. This made it difficult for H4/H5 dimers to form row structures.

### Saturated lipid domains stabilized the row structure of Rh-dimers

Saturated lipid domains were formed due to the affinity of the saturated lipids to raftophilic H8 regions of dimers and the phase separation of saturated and unsaturated lipids (S5 Fig). The stability of the row structures of H1/H8 dimers was influenced by the size of the saturated lipids domains (i.e., the number of saturated lipids in each domain) between two H8 regions of neighboring dimer pairs. The contact between the two Rh-dimers became unstable with decreases in domain size. We determined that ⟨*S*(*t*)⟩ of the model system containing only two H1/H8 dimers decreased with the decrease in the number of saturated lipids (*N*_*s*_) between the two dimers when *N*_*s*_ was less than 5 (Figs 4 and S6). However, ⟨*S*(*t*)⟩_*t*_ for all simulation results (results of ten simulations) exhibited values near the mean when *N*_*s*_ ≥ 6, even though ⟨*S*(*t*)⟩_*t*_ was often near 0 when *N*_*s*_ ≤ 5. This finding indicated that lipid domains containing more than 5 or 6 saturated lipids were essential forming and stabilizing Rh-dimer row structures, not single saturated lipid molecules.

**Fig 4.**
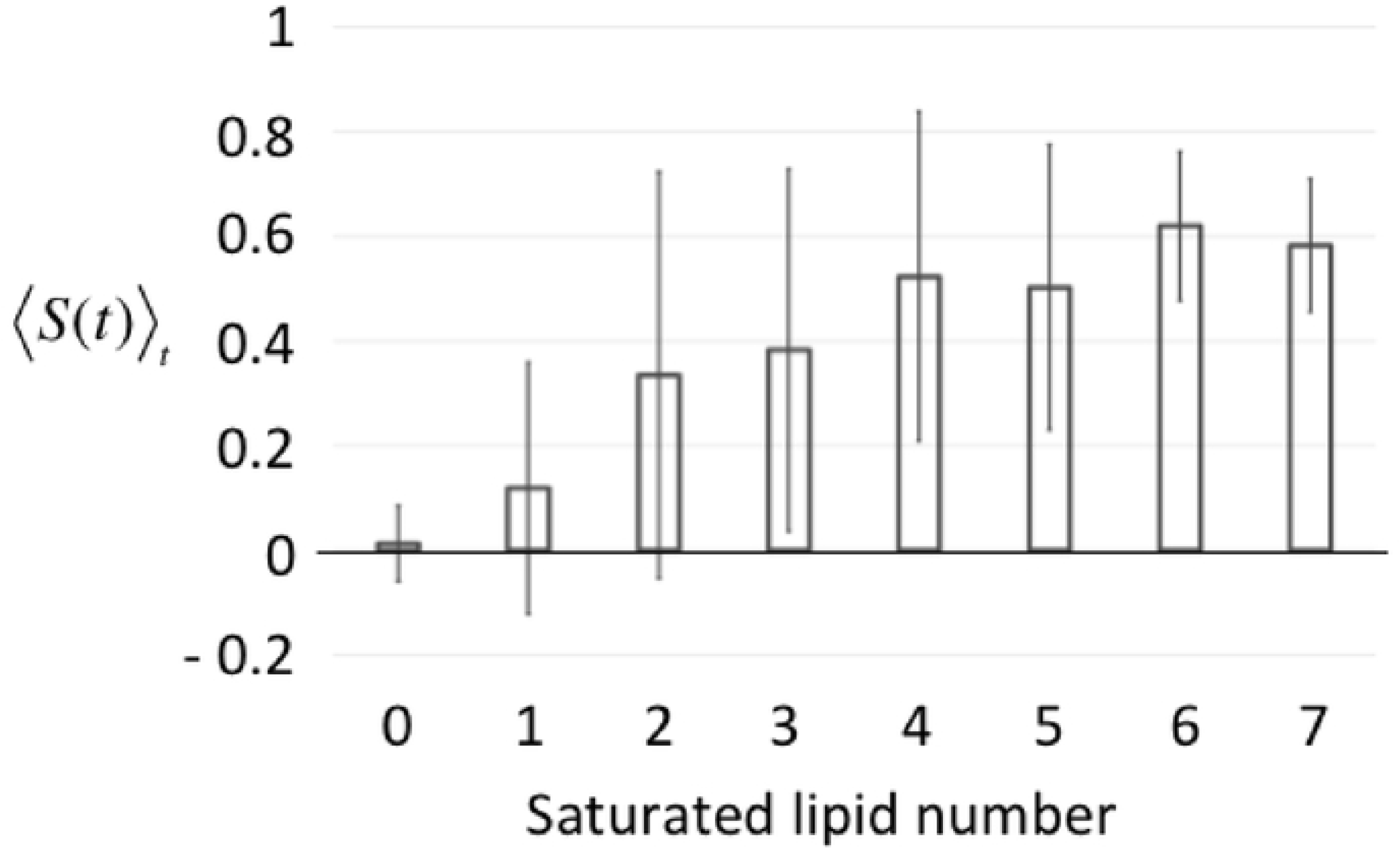
Evaluation of rhodopsin (Rh) H1/H8 dimer low structure stability dependence on saturated lipid domain size. Averages of ⟨*S*(*t*)⟩_*t*_ over ten simulations, ⟨*S*(*t*)⟩ where the system consisted of only two H1/H8 dimers and various numbers of saturated and unsaturated lipids (S5 Fig). Error bars indicate standard deviations. ⟨*S*(*t*)⟩_*t*_ was the average time of *S*(*t*) between 50 to 100 µs and was evaluated according to the change in number of saturated lipids between two Rh-dimers.

## Discussion

In this study, we developed a coarse-grained model of retinal disk membrane consisting of saturated lipids, unsaturated lipids, and Rh-dimers used to simulated row structure formation of Rh-dimers observed in recent experiments. Using this model, we clarified that H1/H8 dimer is the primary element constructing the row structures of Rh dimers. As far as we know, the present study was the first to provide a scenario to explain the mechanism of formation and stabilization of row structure by Rh-dimers in the disk membrane.

Recent experimental evidence have suggested that more than 70% of Rh is in dimeric or higher oligomeric state(s) existing in a dynamic equilibrium [13,18,19]. Based on these findings and our current results, the H1/H8 dimer forms transient row structure through lipid raft-based interaction between the dimers, which appears consistent with results from recent studies [13,31,32]. In addition, we found that the nano-sized domains of saturated lipids with cholesterols (raftophilic lipids, i.e., lipid rafts) were also key factors in forming Rh-dimer row structures with H1/H8 dimers. Such domains acted as “glue” to connect two raftophilic H8 regions of two Rh-dimers to construct the row structures.

In our model, the saturated lipids were assumed to always associate with cholesterols, which allowed the saturated lipids to take rigid structures compared to that of unsaturated lipids. As a result of this assumption, the lipid raft could form by phase separation between saturated and unsaturated lipids. Our simulation strongly suggested that lipid raft formation was essential for the formation of the row of Rh dimes, but did not suggest much about the molecular mechanism of lipid raft formation. For example, weak affinities only among saturated phospholipids could likely confer a similar affect as cholesterol.

In this study, we carefully chose the parameters of the model. However, we found that Rh-dimers could form the row structures if the ratio of rigidity (*q*_*i*_) of saturated and unsaturated lipids and the raftophilicity of Rh-dimers were sufficiently large. This fact provides additional evidence confirming the abovementioned fact that only the H1/H8 dimers and saturated lipid domain formations were essential for Rh-dimer row formation.

The current model was unable to reproduce the repetitions of formation and collapse of row structures of Rh-dimers that have 100 ms range of life time [13,33–35] because of difficulties such as simulation costs for example. The limitation and difficulties will be addressed in future studies by modifying the model to allow for accelerated calculation.

## Acknowledgements

The authors thank H. Nishimori for helpful discussions.

## Author Contributions

Conceived and designed the experiments: AA YK FH KM HY. Performed the experiments: YK AA. Analyzed the data: YK MF AA. Wrote the paper: AA YK MF FH KM HY.

## References

1. Boesze-Battaglia K, Schimmel RJ. Cell membrane lipid composition and distribution: Implications for cell function and lessons learned from photoreceptors and platelets. J Exp Biol. 1997;200: 2927–2936.

2. Miljanich GP, Sklar LA, White DL, Dratz EA. Disaturated and dipolyunsaturated phospholipids in the bovine retinal rod outer segment disk membrane. BBA - Biomembr. 1979;552: 294–306. doi:10.1016/0005-2736(79)90284-0

3. Gahbauer S, Böckmann RA. Membrane-mediated oligomerization of G protein coupled receptors and its implications for GPCR function. Front Physiol. 2016;7: 1–17. doi:10.3389/fphys.2016.00494

4. Levental I, Lingwood D, Grzybek M, Coskun U, Simons K. Palmitoylation regulates raft affinity for the majority of integral raft proteins. Proc Natl Acad Sci. 2010;107: 22050–22054. doi:10.1073/pnas.1016184107

5. Gagnoux-Palacios L, Dans M, van’t Hof W, Mariotti A, Pepe A, Meneguzzi G, et al. Compartmentalization of integrin alpha6beta4 signaling in lipid rafts. J Cell Biol. 2003;162: 1189–96. doi:10.1083/jcb.200305006

6. Hérincs Z, Corset V, Cahuzac N, Furne C, Castellani V, Hueber A-O, et al. DCC association with lipid rafts is required for netrin-1-mediated axon guidance. J Cell Sci. 2005;118: 1687–92. doi:10.1242/jcs.02296

7. Niethammer P, Delling M, Sytnyk V, Dityatev A, Fukami K, Schachner M. Cosignaling of NCAM via lipid rafts and the FGF receptor is required for neuritogenesis. J Cell Biol. 2002;157: 521–32. doi:10.1083/jcb.200109059

8. Shrimpton CN, Borthakur G, Larrucea S, Cruz MA, Dong J-F, López JA. Localization of the adhesion receptor glycoprotein Ib-IX-V complex to lipid rafts is required for platelet adhesion and activation. J Exp Med. 2002;196: 1057–66. doi:10.1084/jem.20020143

9. Zhang W, Trible RP, Samelson LE. LAT palmitoylation: its essential role in membrane microdomain targeting and tyrosine phosphorylation during T cell activation. Immunity. 1998;9: 239–46. doi:10.1016/s1074-7613(00)80606-8

10. Polozova A, Litman BJ. Cholesterol dependent recruitment of di22:6-PC by a G protein-coupled receptor into lateral domains. Biophys J. 2000;79: 2632–2643. doi:10.1016/S0006-3495(00)76502-7

11. Tanimoto Y, Okada K, Hayashi F, Morigaki K. Evaluating the Raftophilicity of Rhodopsin Photoreceptor in a Patterned Model Membrane. Biophys J. 2015;109: 2307–2316. doi:10.1016/j.bpj.2015.10.015

12. Seno K, Hayashi F. Palmitoylation is a prerequisite for dimerization-dependent raftophilicity of rhodopsin. J Biol Chem. 2017;292: 15321–15328. doi:10.1074/jbc.M117.804880

13. Hayashi F, Saito N, Tanimoto Y, Okada K, Morigaki K, Seno K, et al. Raftophilic rhodopsin-clusters offer stochastic platforms for G protein signalling in retinal discs. Commun Biol. 2019;2: 1–12. doi:10.1038/s42003-019-0459-6

14. Najafi M, Haeri M, Knox BE, Schiesser WE, Calvert PD. Impact of signaling microcompartment geometry on GPCR dynamics in live retinal photoreceptors. J Gen Physiol. 2012;140: 249–66. doi:10.1085/jgp.201210818

15. Duft D, Achtzehn T, Müller R, Fotiadis D, Liang Y, Filipek S, et al. Rhodopsin dimers in native disc membranes. Nature. 2003;421: 127–128.

16. Liang Y, Fotiadis D, Filipek S, Saperstein DA, Palczewski K, Engel A. Organization of the G protein-coupled receptors rhodopsin and opsin in native membranes. J Biol Chem. 2003;278: 21655–21662. doi:10.1074/jbc.M302536200

17. Gunkel M, Schöneberg J, Alkhaldi W, Irsen S, Noé F, Kaupp UB, et al. Higher-order architecture of rhodopsin in intact photoreceptors and its implication for phototransduction kinetics. Structure. 2015;23: 628–638. doi:10.1016/j.str.2015.01.015

18. Buzhynskyy N, Salesse C, Scheuring S. Rhodopsin is spatially heterogeneously distributed in rod outer segment disk membranes. J Mol Recognit. 2011;24: 483–489. doi:10.1002/jmr.1086

19. Comar WD, Schubert SM, Jastrzebska B, Palczewski K, Smith AW. Time-resolved fluorescence spectroscopy measures clustering and mobility of a G protein-coupled receptor opsin in live cell membranes. J Am Chem Soc. 2014;136: 8342–8349. doi:10.1021/ja501948w

20. Dell’Orco D, Koch K-W. A dynamic scaffolding mechanism for rhodopsin and transducin interaction in vertebrate vision. Biochem J. 2011;440: 263–71. doi:10.1042/BJ20110871

21. Periole X, Knepp AM, Sakmar TP, Marrink SJ, Huber T. Structural determinants of the supramolecular organization of G protein-coupled receptors in bilayers. J Am Chem Soc. 2012;134: 10959–10965. doi:10.1021/ja303286e

22. Park JH, Scheerer P, Hofmann KP, Choe HW, Ernst OP. Crystal structure of the ligand-free G-protein-coupled receptor opsin. Nature. 2008;454: 183–187. doi:10.1038/nature07063

23. Nagao M, Kelley EG, Ashkar R, Bradbury R, Butler PD. Probing Elastic and Viscous Properties of Phospholipid Bilayers Using Neutron Spin Echo Spectroscopy. J Phys Chem Lett. 2017;8: 4679–4684. doi:10.1021/acs.jpclett.7b01830

24. Levental KR, Lorent JH, Lin X, Skinkle AD, Surma MA, Stockenbojer EA, et al. Polyunsaturated lipids regulate membrane domain stability by tuning membrane order. Biophys J. 2016;110: 1800–1810. doi:10.1016/j.bpj.2016.03.012

25. Salom D, Lodowski DT, Stenkamp RE, Trong IL, Golczak M, Jastrzebska B, et al. Crystal structure of a photoactivated deprotonated intermediate of rhodopsin. Proc Natl Acad Sci. 2006;103: 16123–16128. doi:10.1073/pnas.0608022103

26. Palczewski K. G Protein–Coupled Receptor Rhodopsin. Annu Rev Biochem. 2006;75: 743–767. doi:10.1146/annurev.biochem.75.103004.142743

27. Lin YT, Frömberg D, Huang W, Delivani P, Chacón M, Tolić IM, et al. Pulled Polymer Loops as a Model for the Alignment of Meiotic Chromosomes. Phys Rev Lett. 2015;115: 208102. doi:10.1103/PhysRevLett.115.208102

28. Almeida PFF, Vaz WLC, Thompson TE. Lipid diffusion, free area, and molecular dynamics simulations. Biophys J. 2005;88: 4434–4438. doi:10.1529/biophysj.105.059766

29. Scherfeld D, Kahya N, Schwille P. Lipid Dynamics and Domain Formation in Model Membranes Composed of Ternary Mixtures of Unsaturated and Saturated Phosphatidylcholines and Cholesterol. Biophys J. 2003;85: 3758–3768. doi:10.1016/S0006-3495(03)74791-2

30. Chandrasekhar S. Liquid crystals. 2nd ed. Cambridge: Cambridge University Press; 1992.

31. Knepp AM, Periole X, Marrink SJ, Sakmar TP, Huber T. Rhodopsin forms a dimer with cytoplasmic helix 8 contacts in native membranes. Biochemistry. 2012;51: 1819–1821. doi:10.1021/bi3001598

32. Zhao DY, Pöge M, Morizumi T, Gulati S, Van Eps N, Zhang J, et al. Cryo-EM structure of the native rhodopsin dimer in nanodiscs. J Biol Chem. 2019;294: 14215–14230. doi:10.1074/jbc.RA119.010089

33. Jastrzebska B, Palczewski K, Schubert SM, Smith AW, Comar WD. Time-Resolved Fluorescence Spectroscopy Measures Clustering and Mobility of a G Protein-Coupled Receptor Opsin in Live Cell Membranes. J Am Chem Soc. 2014;136: 8342–8349. doi:10.1021/ja501948w

34. Mishra AK, Gragg M, Stoneman MR, Biener G, Oliver JA, Miszta P, et al. Quaternary structures of opsin in live cells revealed by FRET spectrometry. Biochem J. 2016;473: 3819–3836. doi:10.1042/BCJ20160422

35. Kusumi A, Hyde JS. Spin-label saturation-transfer electron spin resonance detection of transient association of rhodopsin in reconstituted membranes. Biochemistry. 1982;21: 5978–83. doi:10.1021/bi00266a039

